# New 6_3_ knot and other knots in human proteome from AlphaFold predictions

**DOI:** 10.1101/2021.12.30.474018

**Authors:** Agata P. Perlinska, Wanda H. Niemyska, Bartosz A. Gren, Pawel Rubach, Joanna I. Sulkowska

## Abstract

AlphaFold is a new, highly accurate machine learning protein structure prediction method that outperforms other methods. Recently this method was used to predict the structure of 98.5% of human proteins. We analyze here the structure of these AlphaFold-predicted human proteins for the presence of knots. We found that the human proteome contains 65 robustly knotted proteins, including the most complex type of a knot yet reported in proteins. That knot type, denoted **6**_**3**_ in mathematical notation, would necessitate a more complex folding path than any knotted proteins characterized to date. In some cases AlphaFold structure predictions are not highly accurate, which either makes their topology hard to verify or results in topological artifacts. Other structures that we found, which are knotted, potentially knotted, and structures with artifacts (knots) we deposited in a database available at: https://knotprot.cent.uw.edu.pl/alphafold.

## Introduction

The method established by AlphaFold [Jumper et al., 2021] has already led to high-quality predictions for thousands of protein structures from different genomes. AlphaFold’s approach bypasses the need to understand the complex rules that determine the physical protein-folding process. An *in silico* approach can lead to the prediction of new folds and predict structures which are hard to determine using approaches such as crystallization, NMR or cry-EM. Since it is known that the probability that a linear chain is knotted increases with its length, can folding rules be simply bypassed? Are these longer protein structures more complicated than we can currently model accurately?

Recent studies have shown that some proteins form open knots in their native folded structure [Mansfield, 1994, Mallam and Jackson, 2005, Sulkowska, 2020]. These protein knots resemble wellknown rope knots [Sułkowska et al., 2012]. In general, the percentage of known knotted proteins is much lower than would be expected in random polymers with a similar length, compactness, and flexibility [Virnau et al., 2006]. Among known protein structures only the simplest of knot types have been observed [Jamroz et al., 2015] and, further, only ones which can be constructed from one threading [Taylor, 2000]. Interestingly, although their biological role is not clear [Jackson et al., 2017], knotting has been found in all branches of the tree of life and is conserved among sequences with low similarity [Sułkowska et al., 2012]. Their presence raises many fundamental questions, from which we list here just a few: i) Is the small percentage of knotted proteins observed due to the specificity of the data deposited in the PDB? ii) Are there more complex knots which do not yet appear in the PDB? iii) What is the origin of knotted proteins? iv) Is topology strictly conserved? v) What role do knots play in proteins?

Based on human protein structures predicted by AlphaFold, we found answers to some of these questions. We have conducted a comprehensive review of all 23,391 structures predicted by AlphaFold for the human proteome. For each protein structure we determined the dominant knot type [Jamroz et al., 2015] and found the location of the knot cores (i.e. minimal portions of protein backbones that form a given knot type). We found 340 knotted structures (Table 1).However, after a careful evaluation we concluded that over 75% of them are artifacts.In some cases, mostly due to a low level of confidence in crucial parts of the knotted core, we were not able to verify whether or not the structure is knotted. Such structures are labelled *potentially knotted*. Overall, we found 65 robustly knotted structures. These proteins are deposited in an online database to enable further *in vivo* and *in silico* investigation, available at: https://knotprot.cent.uw.edu.pl/alphafold.

**Table 1:** Number of knotted structures with different knot types found within human proteome based on AlphaFold prediction.

| Knot types | No. knotted structures | No. potentially knotted | No. artifact knots |
| --- | --- | --- | --- |
| $3_1$ | 62 | 13 | 179 |
| $4_1$ | 0 | 1 | 12 |
| $5_1$ | 0 | 0 | 10 |
| $5_2$ | 2 | 1 | 16 |
| $6_2$ | 0 | 0 | 3 |
| $6_3$ | 1 | 0 | 0 |
| Other | 0 | 0 | 40 |
| All | 65 | 15 | 260 |

To determine how complex human knotted proteins can be, and whether their topology is conserved, we analyzed the details of these newly identified knotted proteins.

### New type of protein knot found

Thus far four main types of knots have been observed in experimentally solved proteins – 3_1_, 4_1_, 5_2_ and 6_1_ [Jamroz et al., 2015]. Here, we found three types – 3_1_, 5_2_ and 6_3_. The 6_3_ knot has not been seen before in a protein. Note that 4_1_ and 6_1_ knots have not been detected in ourreview, although they have been observed, mostly in bacterial and plant proteins (16 proteins with 4_1_ and two with 6_1_). However, there is one 4_1_ protein (from *Toxoplasma gondii*; UniProtKB ID: B9PJE6) that can potentially have a human counterpart. Similarly, bacterial 6_1_ knotted haloacid dehalogenases [Bölinger et al., 2010] can be present in the human proteome. However, they are not part of the haloacid dehalogenase (HAD) superfamily that groups such proteins, which indicates that they may represent specific types of enzymes. It is plausible that there are no 4_1_ or 6_1_ knots in the human proteome, possibly because the topologies of these proteins were not beneficial enough to be conserved during evolution or because there is simply no need for them in human cells.

### New type of knot – 6_3_ knot

The 6_3_ knot is formed by the backbone of the von Willebrand factor A domain-containing protein 5A (BCSC-1, breast cancer suppressor candidate 1 Figure 1). A knot with six crossings has been found before in haloacid dehalogenases, namely the Stevedore knot (6_1_), which is one of the three prime knots with six crossings (6_1_, 6_2_, 6_3_). Here, for the first time, we observe a second six-crossing knot, 6_3_, in proteins. Moreover, this is the first occurrence of a knot type which is not a twist knot (i.e. is not the result of twists plus one threading [Taylor, 2000]). To form a 6_3_ knot, the protein chain must cross the energy barrier at least twice during folding, as it is pulled through twisted loops.

**Figure 1:**
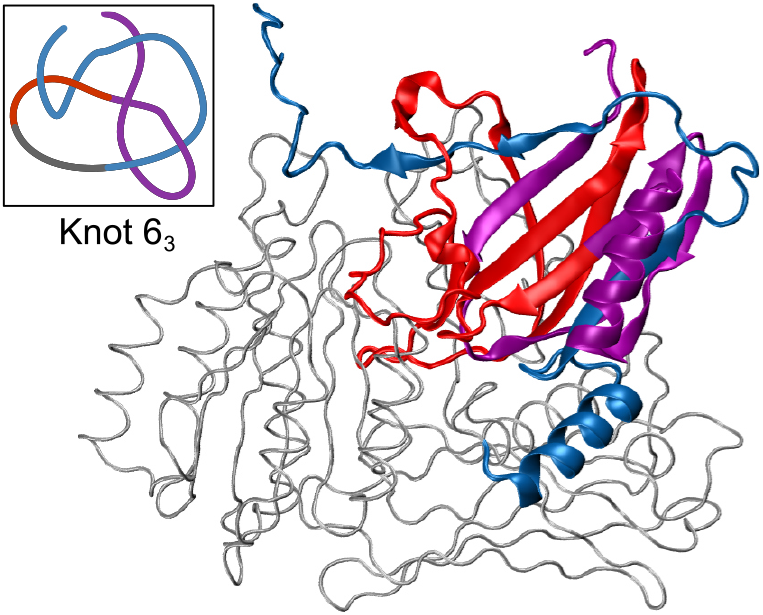
Von Willebrand factor A domain-containing protein 5A (UniProtKB ID: O00534) with a new type of protein knot – 6_3_ shown with essential strands colored in red, blue and purple. Upper left panel shows a simplified knotted core of the protein.

The 6_3_ knot is located in a high confidence region (average per-residue confidence score (pLDDT) is 88.6) between amino acids 45 and 625. It covers most of the protein, which is 786 residues long. However, because it has quite long tails which consist of 45 and 161 amino acids, it is considered to be located rather deep within the structure. This suggests that the knot is stable and will not be untied spontaneously by thermal fluctuations of the protein. Moreover, the presence of the knot is conserved among species (based on rat, mouse, and danio BCSC-1 structures predicted by AlphaFold). However, the danio structure (which is the least similar to the human protein – 57% of sequence similarity) has a 3_1_ knot instead of a 6_3_ knot.

The BCSC-1 protein is interesting not only because of its complex topology, but also because of its crucial function. Despite the lack of information about its structure (or its homologs in other organisms) beforehand, there are studies showing that it is involved in cancer development and can act as its suppressor [Di et al., 2018].

### Known proteins discovered to be knotted

Many of the proteins studied here have experimentally resolved structures. However, the structures often have some unresolved regions (such as loops) or only have one specific domain crystallized. In these cases, determining knotting in the whole chain (and certain subregions) may be impossible. Since AlphaFold predicts structures using their whole sequence, we were able to find knots in proteins that were not considered knotted before.

### Ion channels

Calcium-activated chloride channel regulator 1 (CLCA1; UniProtKB ID: A8K7I4) is a 914 amino acid long protein, only a portion of which was crystallized, domain VWA, 303-459 residues. This domain itself is unknotted since the knotted core is at residues 76-311 (Figure 2B). The knotted core is marked by AlphaFold as a high confidence region (average pLDDT 92.4). Moreover, we found that its paralogs (also predicted by AlphaFold: CLCA2 and CLCA4) have the same knot type and location.

**Figure 2:**
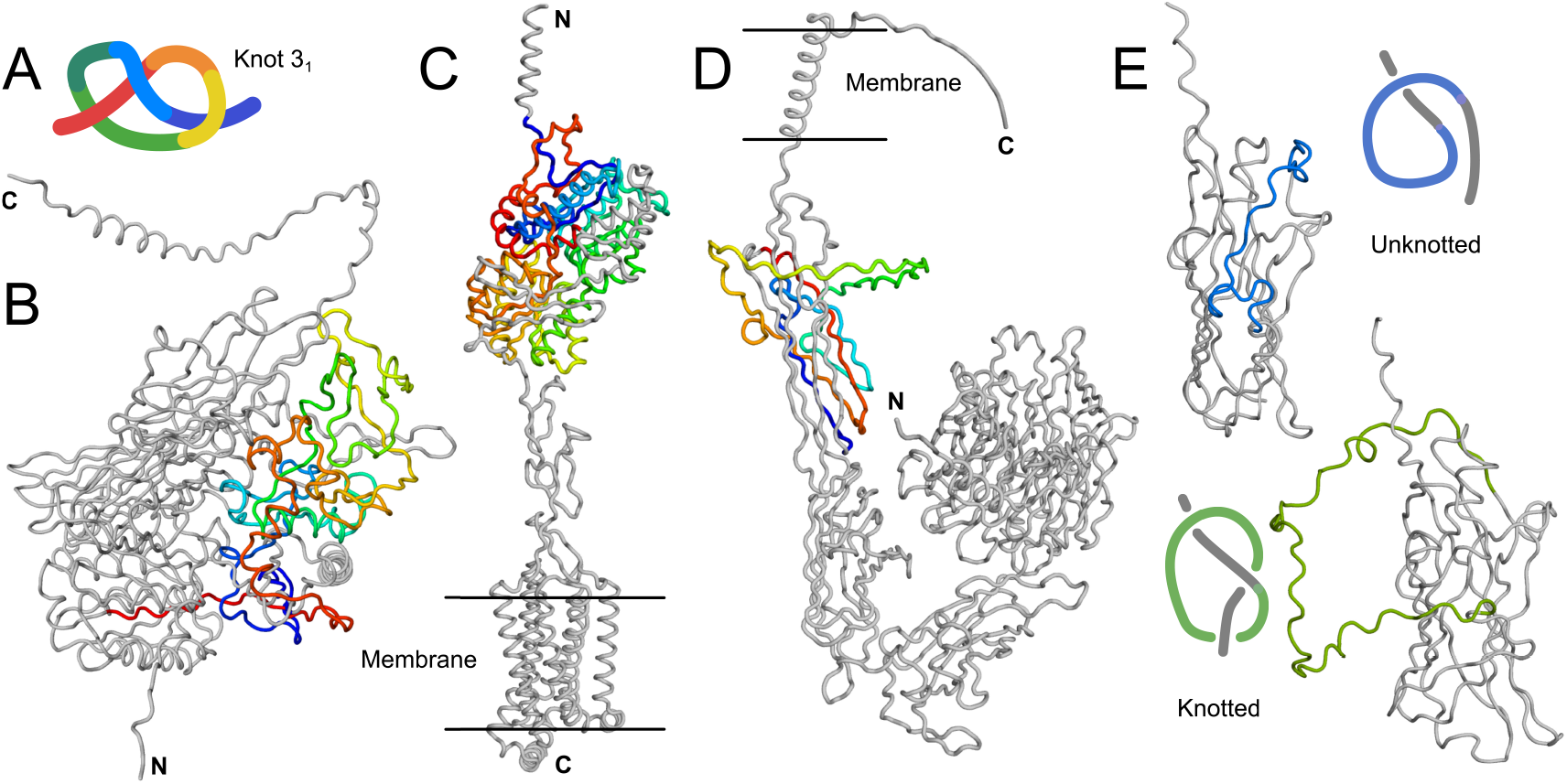
Selected proteins with a 3_1_ knot. (A) Simplified representation of a 3_1_ knot found in proteins. (B) Calcium-activated chloride channel regulator 1 (UniProtKB ID: A8K7I4). (C) Taste receptor type 1 member 2 (UniProtKB ID: Q8TE23). (D) Integrin alpha IIb (UniProtKB ID: P08514). All knotted cores are shown in rainbow color scheme to guide the eye. (E) Upper panel: unknotted integrin alpha 1 (UniProtKB ID: P56199). Lower panel: 3_1_ knotted integrin alpha 3 (UniProtKB ID: P26006).

CLCA proteins undergo self-cleavage (CLCA1 at position 695) to form the N- and C-terminal portions of the proteins. The N-terminal portion is responsible for the function of the protein, activating ion channels [Yurtsever et al., 2012]. The fact that the N-terminal portion is knotted was unknown until now, which emphasizes the importance of having full structure information available.

### Taste receptors

Taste receptor type 1 member 2 (TAS1R2; UniProtKB ID: Q8TE23) is another example of a 3_1_ knotted protein. While the structure of this human protein is not yet resolved experimentally, its orthologs from other organisms are (e.g. *Oryzias latipes* – Japanese rice fish; UniProtKB ID: A0A173M0G2; 34% of sequential identity). Interestingly, knots have not been detected in any of the orthologs because of the gaps in the sequence combined with truncated data at the N-terminal end.

The knotted core in the protein is located at residues 20-389 and encompasses most of its extracellular part (Figure 2C). TAS1R2, together with TAS1R3 (also knotted as per AlphaFold prediction), makes a heterodimeric transmembrane sweet taste receptor. It is responsible for the perception of natural sugars (also artificial sweeteners) with its extracellular domain where they bind.

### Knot or not?

Often experimentally resolved structures possess missing regions that are not structurally organized. Such regions can be crucial for making the knot since the way they will be implemented into the structure can make a knot… or not. The integrins alpha described below underline the importance of this aspect.

### Integrins alpha

We found the 3_1_ knot in 7 out of 18 human integrins alpha. In each of them, the knot is located in the C-terminal part of the integrin (in domain called Calf2; Figure 2D). Based on the AlphaFold predicted structures, it appears that whether the integrin will be knotted is determined by a single loop. Only integrins with longer loops are knotted (green region in Figure 2E). Unfortunately, this loop in most of the integrin alpha structures has a low confidence (pLDDT) score making its location uncertain. Therefore, we cannot verify whether or not the knot present in these proteins is an artifact. If some of the integrins are indeed knotted, they would provide a second example of a family containing topologically distinct proteins – the first being aspartate/ornithine carbamoyltransferases [Sulkowska, 2020].

### Artifacts

More than 75% of knotted human structures from AlphaFold we determined as artifacts. In such structures the knot is often created in low confidence regions (pLDDT ¡ 50), which means its position is not reliable. Moreover, about 1/3 of the structures with artifact knots represent large proteins (*>* 2, 700 aa) that were predicted using multiple overlapping models. In fact, more than 50% of the artifact structures, yet only 17% of the robustly knotted structures, have more than 1,000 residues. Since it is increasingly difficult to correctly predict structure and topology with increasing protein length and without known type of knots, topology can be used as a tool to identify the quality of experimental or simulated data. The examples of interesting artifacts are described on the website.

## Conclusion

While knots are relatively rare in proteins, their conservation suggests a mysterious functional utility. Analyses of entanglement can help to validate structures and provide opportunities to find the interpretation of experimental and simulation data, as well as to provide targets for further studies. Specifically, it would be beneficial to experimentally verify several proteins, including those of uncertain topology, such as chloride channel protein 2 (UniProtKB ID: P51788). If the detected 5_2_ knot truly exists, it would be a second type of proteins with this complex knot. Furthermore, this data, as well as structures predicted for other proteomes, can help to understand how evolution coded higher levels of protein organization, such as the non-trivial topology.

## Materials and methods

The human proteome was downloaded from the AlphaFold Protein Structure Database [Jumper et al., 2021]. It consisted of 23,391 structures (20,504 proteins). All of the structures were analyzed by computing the HOMFLY-PT polynomial for 200 random closures. The details of the method are explained in [Jamroz et al., 2015]. We determined the position of the knot core in the structure based on the so-called matrix model [Jamroz et al., 2015]. The structure is classified as robustly knotted when random closures form a nontrivial knot more frequently than a trivial knot, the pLDDT score for the knot core is above 50%, and no suspicious geometry was found after a visual inspection. The structures which did not pass the visual inspection are called potentially knotted. The structures with obvious problems are called artifacts.

## Acknowledgements

We thank you Eric Rawdon and Andrzej Stasiak for proof reading this article. This work was supported by EMBO Installation Grant 2057, the Polish Ministry for Science and Higher Education 0003/ID3/2016/64 (to JIS) and supported by COST EUTOPIA action.

